# Deep phylotaxonogenomic revision of the genus *Xanthomonas* within family *Xanthomonadaceae* and proposal of novel family *Frateuriaceae* within order *Xanthomonadales*

**DOI:** 10.1101/2022.09.22.508540

**Authors:** Kanika Bansal, Sanjeet Kumar, Anu Singh, Arushi Chaudhary, Prabhu B. Patil

## Abstract

Genus *Xanthomonas* is primarily comprise phytopathogenic species. In a recent study by carrying out deep phyto-taxonogenomics, we reported that even the genera *Xylella, Stenotrophomonas* and *Pseudoxanthomonas* are miss-classified and belong to genus *Xanthomonas*. Hence to understand the breadth of the genus, we carried out deep phylo-taxonogenomics of the order *Xanthomonadales*. Such investigation revealed that at least four more genera belong to genus *Xanthomonas* with prominent being *Lysobacter*. Further order level deep phylo-taxonogenomics revealed two major families. One being the original family *Xanthomonadaceae* and other is proposed as *Frateuriaceae* fam. nov. as synonym of family *Rhodanobacteraceae* with novel genus *Frateuria* gen. nov.

## Introduction

According to the current standing in nomenclature order *Lysobacterales* (earlier described as *Xanthomonadales*) constitutes two families, namely: *Lysobacteraceae* (earlier described as *Xanthomonadaceae*) and *Rhodanobacteraceae* (LPSN). Both these complex families of the order consist of more than 30 genera and around 300 species [1]. These include diverse species of phytopathogens, environmental pathogens, and opportunistic human pathogens [2]. The *Lysobacteraceae* family contains several species of major plant pathogens with significant economic and agricultural impacts [3-6]. Species of the genus *Xanthomonas* and *Xylella* are phytopathogenic in nature, infecting a wide range of agricultural crops and economic plants. On the other hand, the family also includes the genus *Stenotrophomonas*, which is a multidrug-resistant opportunistic pathogen, responsible for hospital-acquired infections in immunocompromised patients. Environmental members of this family include genera *Pseudomarimonas, Lysobacter*, and *Vulcaniibacterium* etc. which includes species of industrial importance with the ability to cause leaching of heavy metals and bioremediation [7-9]. On the other hand, the family *Rhodanobacteraceae* [1], consists of genera encoding functions required for bioremediation of hydrocarbons, etc. [10, 11].

Integration of classical taxonomy with genomic evidences has revolutionized the field of microbial genomics [12]. Implementation of genome-based methods has enabled several reconciliations at the order, family, or species level. Average nucleotide identity (ANI) is widely used for species delineation; however, it is not suitable for genus-level taxonomy. Average amino acid identity (AAI) and percentage of conserved proteins (POCP) have been proposed for genus-level classification [13, 14]. Out of these, POCP only accounts for the presence or absence of protein. However, AAI is based on identity calculation of overall protein sequences. Hence, AAI is more robust toward taxonomy at the genus level [15]. Though the genus level threshold for AAI was assigned to be 60%, Meehan and co-workers have used 65% and Zheng *et al*. used 68% cutoffs for the genus delineation of the genus *Mycobacterium* and *Lactobacillus*, respectively [15, 16]. Since AAI values may also be affected by the lateral gene transfer. Hence, AAI values for the core genes (cAAI) are also nowadays used as a robust parameter for genus delineation [15, 16]. In addition to these genome similarity criteria, whole genome-based phylogeny provides us with a robust phylogenomic framework, which is crucial evidence for taxonomy.

The current taxonomy of the order is based on 16S rRNA, molecular markers, conserved sequence indels (CSI) and phylogenomics based on no more than 30 conserved proteins [1]. The first whole-genome information-based study of the order provided by our group with the major reshufflings at the family level apart from revealing the boundary of the order and its outliers [7]. In a recent study, by carrying out a deep phylo-taxonogenomic investigation of *Xanthomonas* with its close relatives, we reported that members of genera *Pseudoxanthomonas, Stenotrophomonas*, and *Xylella* belong to a single genus (Bansal et al.,2020). As an extension of the previous study, in the present study we re-evaluate the family and genus boundaries of the order by including all the type species or representative species of the member species of order the *Lysobacterales* (*Xanthomonadales*). We have extended our study to evaluate the taxonogenomic and phylogenomic relationships amongst the genera and provide the description of order *Xanthomonadales* and family *Xanthomonadaceae* as earlier synonyms of order *Lysobacterales* and family *Lysobacteraceae*. Also, we propose a novel genus *Fratura*, and family *Fraturacea* fam. nov. within order *Xanthomonadales*.

## Materials and Methods

### Genome procurement and genome quality assessment

All the information of member within the order the *Xanthomonadales* was obtained from their LPSN definition (https://lpsn.dsmz.de/order/lysobacterales). Genomes of the strains used in the study was obtained from NCBI microbes (https://www.ncbi.nlm.nih.gov/genome/microbes/). The quality checking of the genomes was carried out using checkM v1.1.3 [17]. The genomes that passed the QC (< 5% contamination and < 5% contamination) were considered and annotated using prokka v1.14.6 [18].

### Phylogenomics investigation of the member species of the order *Xanthomonadales*

Core genome phylogeny was generated using PhyloPhAn which is based on more than 400 conserved genes [19]. Here, USEARCH v5.2.32 [20] was implemented for ortholog searching, MUSCLE v3.8.3 [21] for multiple sequence alignment and FastTree v2.1 [22] for phylogenetic construction were used. In order to obtain, a more robust phylogeny, we have fetched core gene using PIRATE [23]. PIRATE is suitable for the identification of orthologue groups of divergent genomes by using amino acid identity thresholds of 50%, 60%, 70%, 80%, 90% and 95%. PIRATE executes a pangenome pipeline to fetch the core genome with a high level of robustness.

### Taxonogenomic assessment of the member species of order *Xanthomonadales*

Genome relatedness amongst the genomes was assessed using average amino acid identity (AAI) using CompareM v0.0.23 (https://github.com/dparks1134/CompareM). Further, for core average amino acid identity (cAAI), core genes amongst the type/representative species genomes were fetched from the PIRATES pangenome analysis. These core genes were then used to evaluate the cAAI values.

## Results and discussion

### Genomic features of order the *Lysobacterales* (*Xanthomonadales*)

A total of 213 genomes (above the quality control of completeness and contamination) belonging to type species and type strains of the order *Xanthomonadales* are summarised in table (Supplementary Table 3). Species of genus the *Xanthomonas* have a genome size of 5 Mb, whereas *Pseudoxanthomonas* and *Stenotrophomonas* have 3–5 Mb genomes, and *Xylella*’s genome is 2.5 Mb in size. Other members of the order, such as *Lysobacter, Luteimonas, Thermomonas*, and *Vulcaniibacterium* have genomes ranging in size from 2-4 Mb. The average GC content of species of genus *Xanthomonas, Pseudoxanthomonas*, and *Stenotrophomonas* genomes has an average GC content of 60–70%, whereas species of genus *Xylella* have a reduced GC of around 51%. Other species of the genus have a 65-71% GC content. In contrast, other species of the remaining genus, such as *Thermomonas, Rhodanobacter, Frateuria, Dyella, Luteibacter* etc. have 2.5-5 Mb genomes with 60-65% GC.

### Phylogenomic evaluation of the order *Lysobacterales* (*Xanthomonadales*)

A total of 213 genomes (above the quality control of completeness and contamination) belonging to type species and type strains of the order *Xanthomonadales* are summarised in table (Supplementary Table 3). A whole genome-based phylogeny using PhyloPhlAn including the type species of the order *Lysobacterales* (n=33), primarily suggest two major clades corresponding to the families of *Lysobacteraceae* (*Xanthomonadaceae*) and *Rhodanobacteraceae*, which we have labelled as *Frateuriaceae* (Pl see later sections) (Figure 1). *Pseudofulvimonas gallinarii* and *Ahniella affigens* formed outgroup for these families respectively. A whole genome-based phylogeny including all the members species (type strains and representative strains) of the order *Lysobacterales* also revealed two major clades, representing families of *Xanthomonadaceae* and *Rhodanobacteraceae*, which we have labelled as *Frateuriaceae* (Pl see later sections) (Figure 2). PhyloPhlAn uses more than 400 conserved genes across bacterial systems suggesting a robust phylogenomic positioning of the species. The phylogenomic positioning of the constituent species was further confirmed using a core genome-based tree. Implementation of PIRATE (pan genome investigation) resulted in a core genome extraction from all the strains (n=213). A phylogenomic tree obtained using FastTree resulted in similar phylogenomic tree as obtained using PhyloPhlAn (Figure 2, Figure 3). In all the three phylogenomic tree construction, we used *Ignatzshineria larvae* DSM 13226^T^ and *Pseudomonas aeruginosa* DSM50071^T^ as the outgroup.

**Figure 1:**
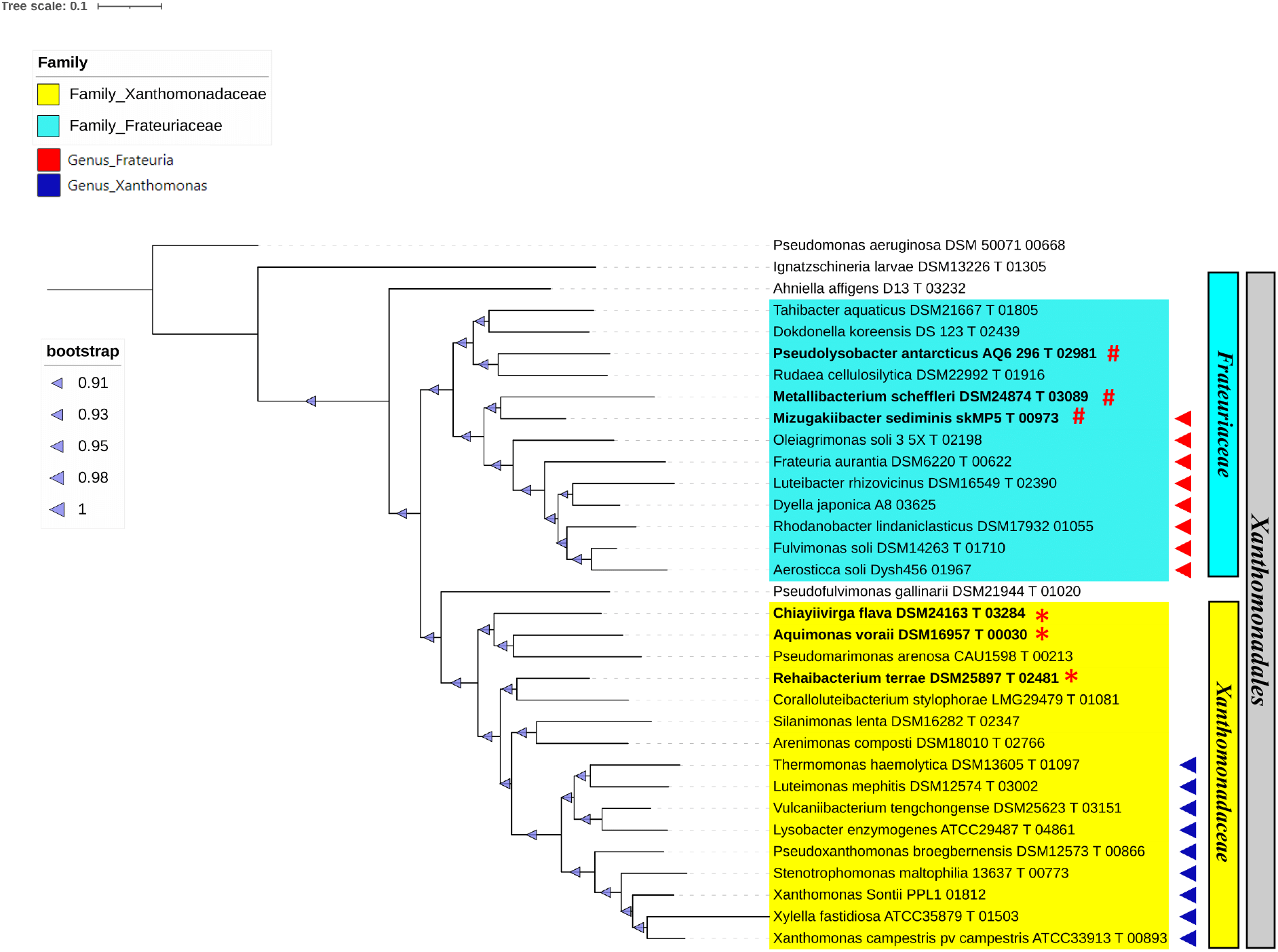
PhyloPhlAn of type species. PhyloPhlAn tree comprising 33 type species and two outgroups, *Pseudomonas aeruginosa* DSM 50071 00668 and *Ignatzschineria larvae* DSM13226 T 01305. Yellow represents the *Xanthomonadaceae* family, with blue triangles representing species in the genus *Xanthomonas*. The sky-blue tint represents the *Frateuriaceae* family, while the red triangles represent the *Frateuria* genus. On the left side, the color legends and bootstrap values are displayed.

**Figure 2:**
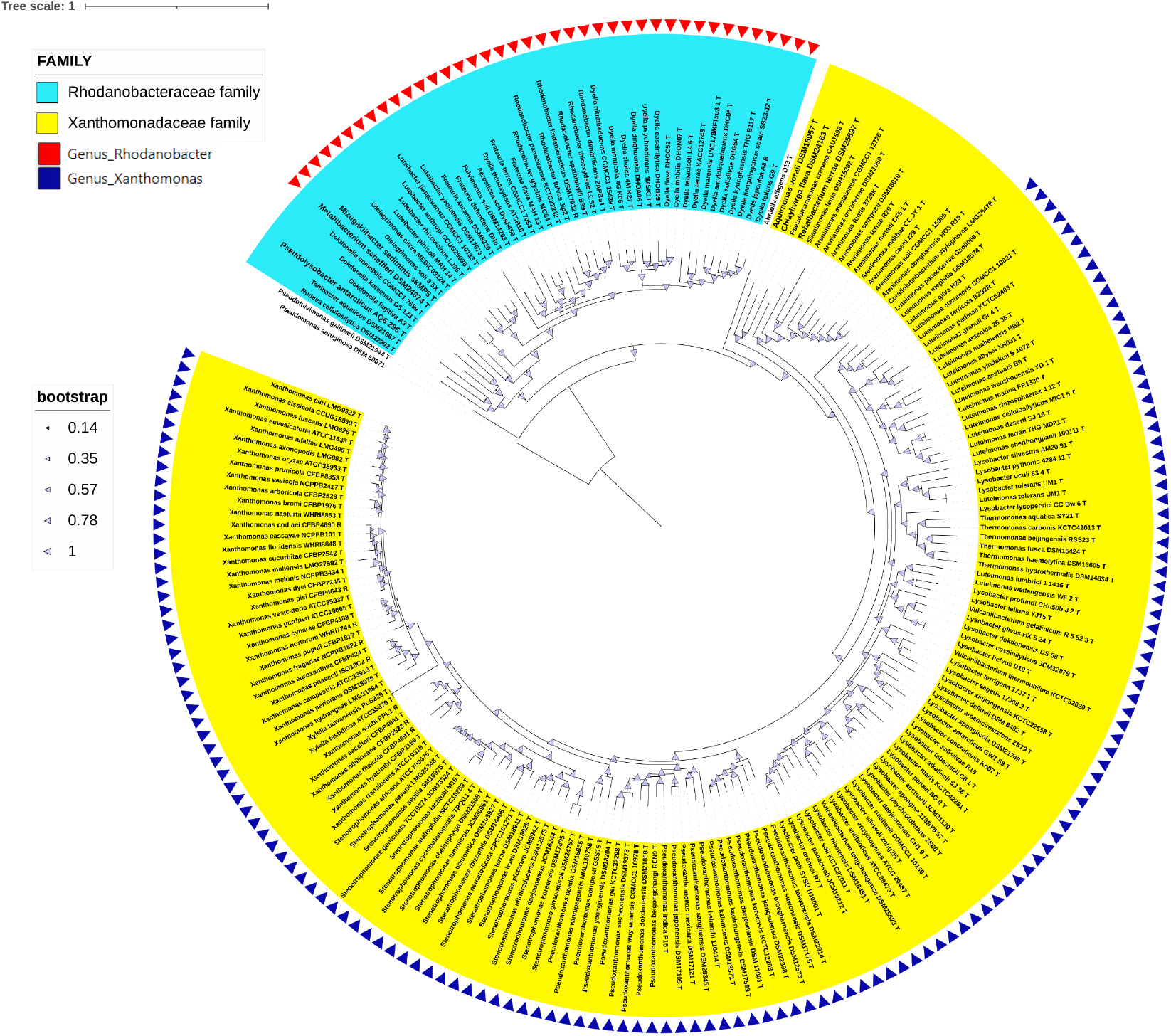
PhyloPhlAn of type strains. PhyloPhlAn tree depicting 213 Type strains, with *Pseudomonas aeruginosa* DSM 50071 00668 serving as an outgroup. Yellow shade represents the *Xanthomonadaceae* family, with blue color triangles representing species belonging to the genus *Xanthomonas*. The *Frateuriaceae* family is represented by the sky-blue hue, while the genus *Frateuria* is represented by the red triangles. On the left side, the color legends and bootstrap values are displayed.

**Figure 3:**
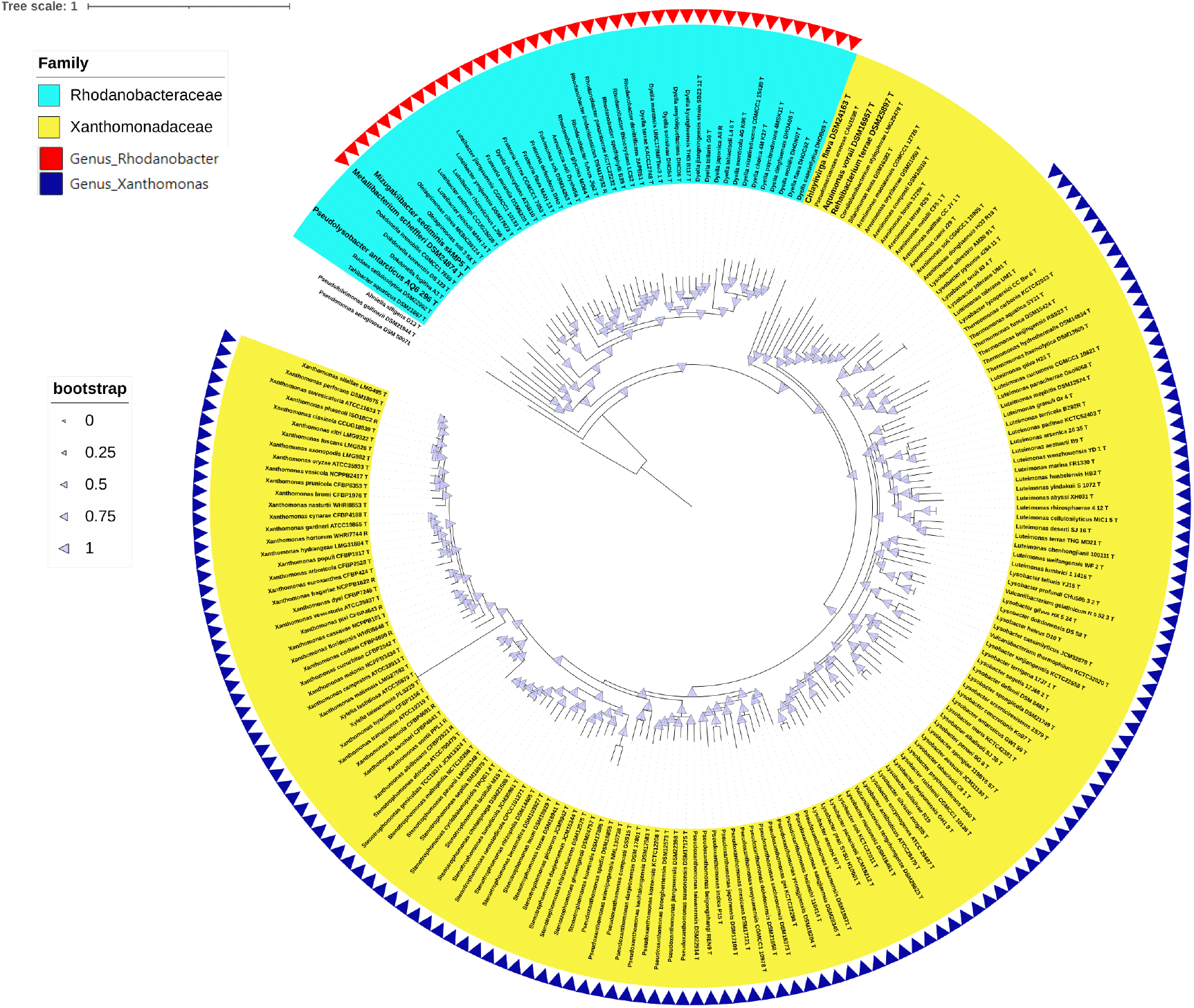
PIRATE of type strains. Core-genome based PIRATE tree is shown with 213 Type strains and *Pseudomonas aeruginosa* DSM 50071 00668 as outgroup. Yellow hue represents the *Xanthomonadaceae* family, with blue color triangles representing species belonging to the genus *Xanthomonas*. The *Frateuriaceae* family is represented by the sky-blue hue, while the genus *Frateuria* is represented by the red triangles. On the left side, the color legends and bootstrap values are indicated.

**Figure 4:**
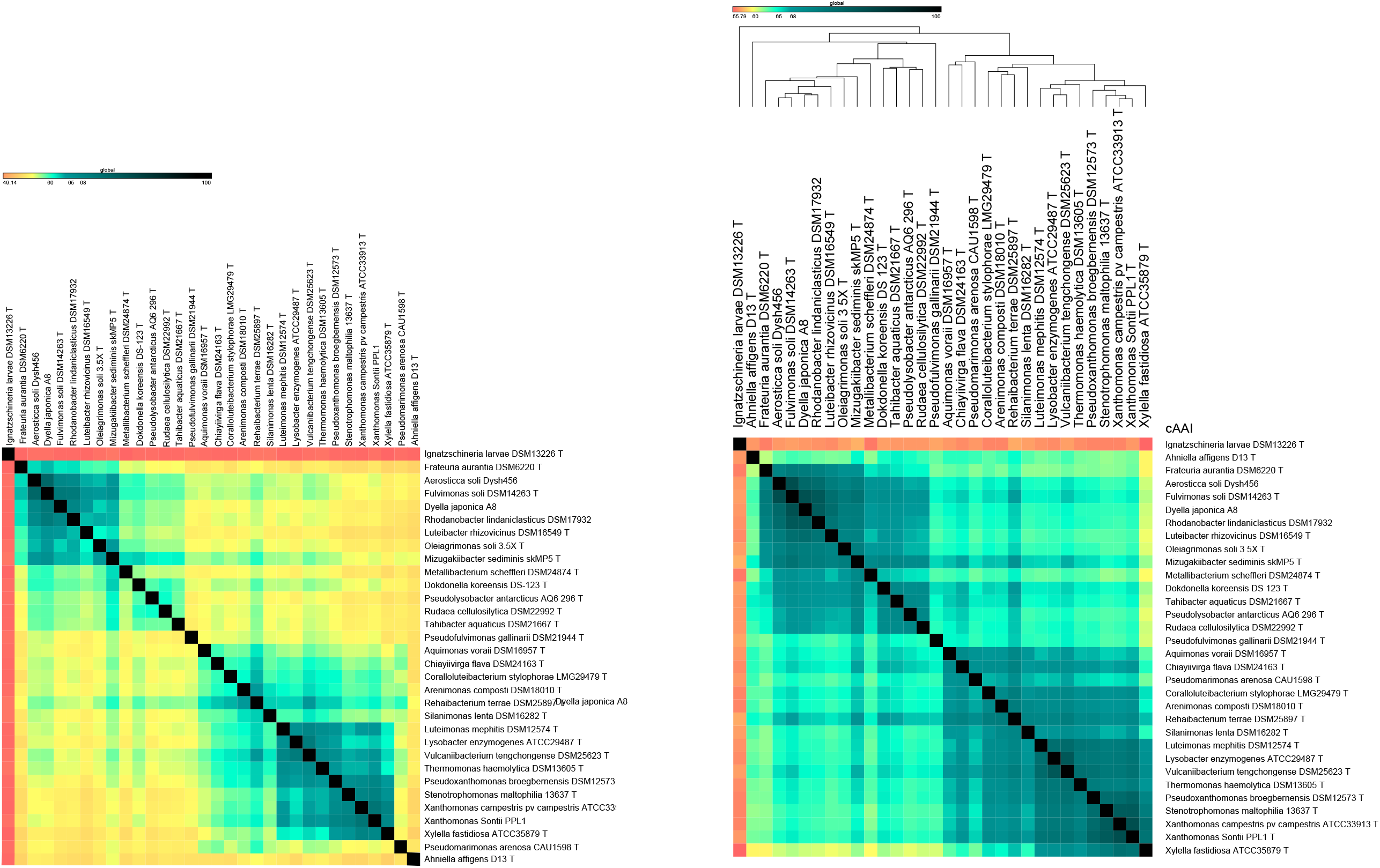
AAI and cAAI of type species. Heatmap showing the average amino acid identity (AAI) and core average amino acid identity (cAAI) of 33 type species. The sky-blue color boxes symbolize the *Xanthomonadaceae* and the *Frateuriaceae* family.

**Figure 5:**
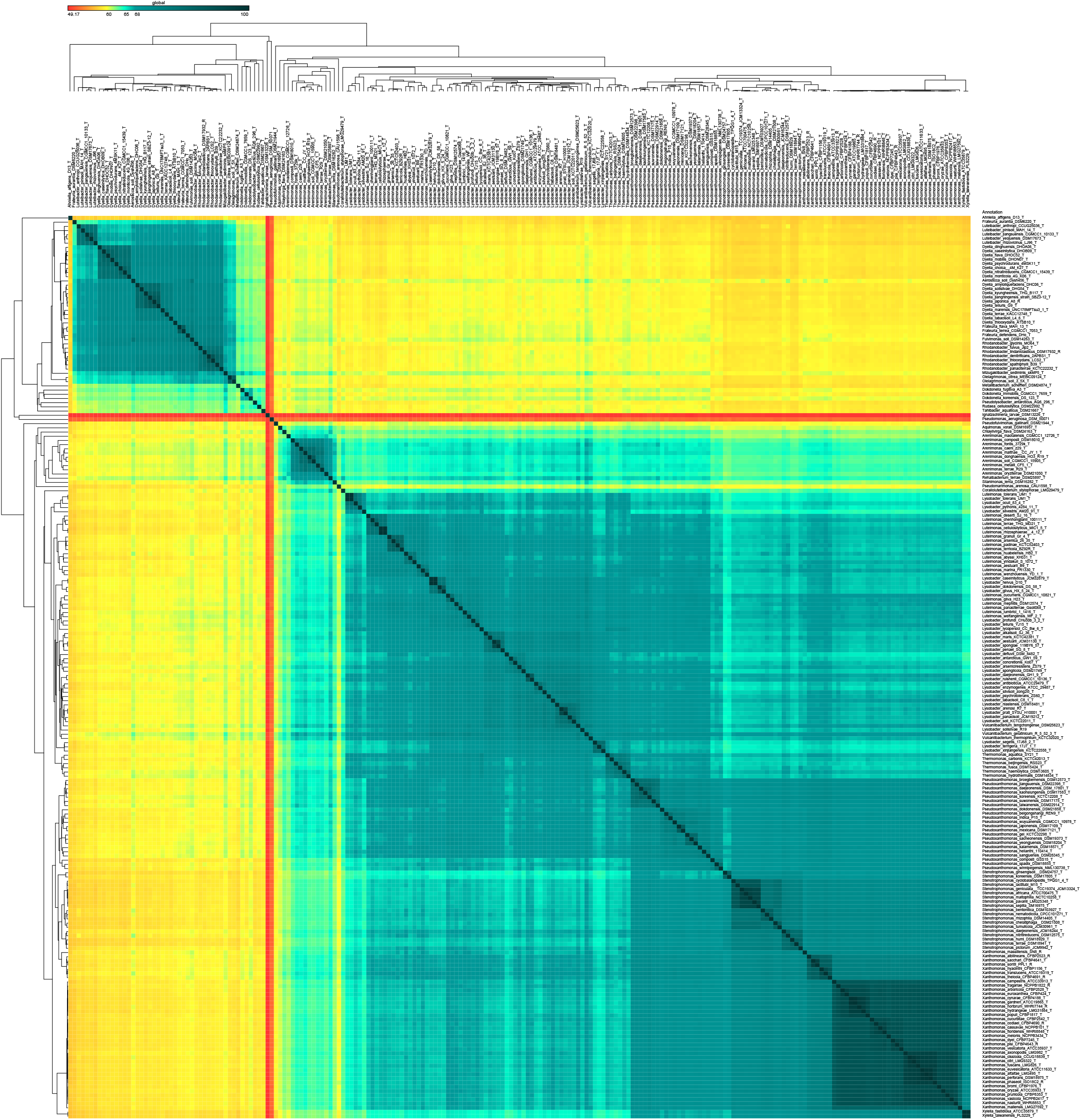
AAI of type strains. Heatmap showing average amino acid identity (AAI) values amongst 213 type strains. The *Xanthomonadaceae* and the *Frateuriaceae* families are represented by the sky-blue boxes.

Here, genera which according to current standing in nomenclature, are known to belong to the families *Lysobacteraceae* and *Rhodanobacteraceae* were found to be correct according to the core genome phylogenies in the present study, with few an exceptions of genus reshufflings across two families.

A genome-based phylogeny of the order *Lysobacterales* was carried out using type/representative species of each genus (Figure 1) as well as type/representative strains of each genus (Figures 2 and 3). Two alternative methods of phylogeny analysis were used in the current study. The phylogeny in the first method was created using more than 400 core genes that are conserved in the bacterial domain (Figure 1 and 2). While, in another approach, the core genes were fetched from the order *Lysobacterales* by using a pan-genome pipeline (Figure 3). Here, *Ignatzshineria larvae* DSM 13226 (T) and *Pseudomonas aeruginosa* DSM50071 (T) were used as the outgroups for the phylogeny. A pangenome investigation was performed to obtain pan genes and core genes of the order *Lysobacterales*, including outgroups *Ignatzshineria larvae* DSM 13226 (T) and *Pseudomonas aeruginosa* DSM50071 (T). In the core genes-based trees, the two families *Lysobacteraceae* and *Rhodanobacteraceae*, which we have labelled as *Frateuriaceae* (Pl see later sections) are highlighted in the figures. Here, genera *Aquimonas, Chiayiivirga*, and *Rehaibacterium* were in the phylogroup of the family *Lysobacteraceae* while, genera *Mizugakiibacter, Metallibacterium*, and *Pseudolysobacter* were in the phylogroup of the family *Rhodanobacteraceae* (Figure 1, 2 and 3).

### Taxonogenomic assessment of genus of *Xanthomonas* and *Rhodanobacter* within their respective families

These new approaches to phylogeny have eliminated some of the biases in previous methods that might have led to incorrect interpretation of phylogenetic data. Based on whole-genome phylogeny, the previously described genus within the order *Xanthomonadales*, such as *Pseudofulvimonas* and *Ahniella* suggests a phylogenetic positioning outside their respective families. However, pairwise comparison of the nearly complete 16S rRNA gene sequence was lower than 93% with the species of order *Xanthomonadales*. The taxonomic boundaries based pairwise sequence similarity of the 16S rRNA gene sequence across the genera, families and orders as around and lower than 94.5%, 86.5%, and 82% respectively [23].Furthermore, to determine the taxonomic status of the genera belonging to the order *Lysobacterales* (*Xanthomonadales*), we have performed overall genomic relatedness index (OGRI) along with taxonogenomic analysis for all the type/representative species and for all type/representative strains of the order. Here, according to the Average Amino acid Identity (AAI) and core Average Amino acid Identity (cAAI) cut-off values of 65% used for delineating novel genus, amongst type/representative species, the genus boundary for *Xanthomonas* extends by including eight genera. Three of these genera were already reported to be synonyms in an earlier in-depth phylo-taxonogenomic study [24]. Here, according to the genus cut-off, all these eight genera are synonyms of genus *Xanthomonas* (Figure 1, Figure 2). These results also correlate with the whole genome-based phylogeny, as both this genus form a distinct phylogroup within the family (representative by the bold font in the phylogenomic tree) (Figure 1, 2 and 3). In this context, the classification of genus within *Xanthomonas* warrants reconstitution and the need for reapplication of origin genus and family label i.e., *Xanthomonadaceae* that we are proposing in the present study. Similarly, according to genus level cut-off for AAI and cAAI, the genus *Rhodanobacter* includes seven more genera: *Aerosticca, Fulvimonas, Dyella, Luteibacter, Frateuria, Oleiagrimonas* and *Mizugakiibacter* that correspond to another major phylogroup within family *Rhodanobacteracea*. With these seven genera being synonyms and the genus *Frateuria* having precedence over other genera and we are proposing family *Frateuriaceae* fam.nov. Since, the genus *Lysobacter* is a synonym of *Xanthomonas* which makes the proposal of order *Xanthomonadales* and family *Xanthomonadaceae* by Saddler and Bradbury 2005 (a, b) [2, 25] is not illegitimate [1].

Based on these results, we are proposing the following emended description of order *Xanthomonadales*, description of family *Xanthomonadaceae*, description of family *Frateuriaceae*, description of family *Pseudofulvimonadaceae*, emended description of genus *Xanthomonas* and its constituent member species, description of genus *Frateuria* and its constituent member species, description of genus *Aquimonas*, description of genus *Chiayiivirga*, genus *Rehaibacterium*, description of *Pseudolysobacter*.

### Emended description of the order *Xanthomonadales* [Saddler and Bradbury 2005 a, b) [2, 25]

#### Synonym: *Lysobacterales* Christensen and Cook (1978) (Approved Lists 1980)

The order consists of two families, *Xanthomonadaceae* and *Frateuriaceae*. The characteristics of the organisms in the order is as described by Naushad et al., 2014 [1] and Saddler and Bradbury (2005 a, b) [2, 25] and based on phylo-taxonogenomic analysis in the present. The type genus is *Xanthomonas* (*Xanthomonas campestris* (Pammel 1895) Dowson 1939 (Approved Lists 1980)) as *Lysobacter* (*Lysobacter* Christensen and Cook 1978 (Approved Lists 1980)) is re-classified as a synonym of *Xanthomonas* in the present study.

#### Emended description of the family *Xanthomonadaceae* Saddler *et. al* 2005

N.L. fem. n. *Xanthomonas*, type genus of the family; L. fem. pl. n. suff. -aceae, ending to denote a family; N.L. fem. pl. n. *Xanthomonadaceae*, the *Xanthomonas* family.

Synonym: *Lysobacterales* Christensen and Cook 1978 (Approved Lists 1980)

The family is within the order *Xanthomonadales* and includes the genera *Xanthomonas* Dowson 1939 [26], *Coralloluteibacterium* Chen et al. 2018 [27], *Arenimonas* Kwon et al. 2007 [28], *Silanimonas* Lee et al. 2005 [29], *Rudaea* Weon et al. 2009 [30], *Chiayiivirga* Hsu et al. 2013 [31], *Aquimonas* Saha et al. 2005 [32], *Pseudomarimonas* Weerawongwiwat et al. 2021 [33]. The description of the family *Xanthomonadaceae* is as given by Saddler *et. al* 2005 and Christensen and Cook 1978 (Approved Lists 1980) with the following amendments, of inclusion in *Aquimonas* Saha et al. 2005 [32], *Chiayiivirga* Hsu et al. 2013 [31], *Rehaibacterium* Yu et al. 2013 [34] and exclusion of *Mizugakiibacter* Kojima et al. 2014 [35], *Metallibacterium* Ziegler et al. 2013 [36] and *Pseudolysobacter* Wei et al. 2020 [37]. The member genera of the family have been established by the latest deep phylotaxonogenomic analysis in the present study.

### Description of the family *Frateuriaceae* fam. nov

*Frateuriaceae* (Frat.eur’i.a.a,ce’ae N.L. fem. n. *Frateuria* is the type genus of the family; -aceae represents the family; N.L. fem. pl. n. *Frateuriaceae* the family whose nomenclature type is the genus *Frateuria*).

Synonym: *Rhodanobacteraceae* Naushad *et al*. 2015 [1]. The family is within the order *Xanthomonadales*. The proposed genus including *Frateuriaceae* includes the genus *Frateuria* Swings et al. 1980 [38], *Dokdonella* Yoon et al. 2006 [39], *Pseudolysobacter* Wei et al. 2020 [37], *Rudaea* Weon et al. 2009 [30], *Tahibacter* Makk et al. 2014 [40]. Exclusion in *Aquimonas* Saha et al. 2005 [32], *Chiayiivirga* Hsu et al. 2013 [31], *Rehaibacterium* Yu et al. 2013 [41] from previously described family *Rhodanobacteraceae* and inclusion of *Mizugakiibacter* Kojima et al. 2014 [35], *Metallibacterium* Ziegler et al. 2013 [36] and *Pseudolysobacter* Wei et al. 2020 [37]. The member genera of the family have been identified by the latest deep phylotaxonogenomic analysis in the present study.

### Description of the family *Pseudofulvimonadaceae* fam. nov

N.L. fem. n. *Pseudofulvimonas*, type genus of the family; L.fem. pl.n, suff. -aceae, ending to denote a family; N.L. fem. Pl. n. *Pseudofulvimonadaceae*. The family includes the genus *Pseudofulvimonas* and *Ahniella* which were identified based on deep phylotaxonogenomic analysis in the present study.

### Emended description of the genus *Aquimonas* Saha et al. 2005

*Aquimonas* (A.qui.mo.nas. L. fem. n. aqua, water; L. fem. n. monas, a unit, monad; N.L. fem. n. *Aquimonas*, a water monad, referring to the isolation of the type species from a warm spring water sample). The type species is *Aquimonas voraii* [32] and it is on the approved list of 1980. The characteristics of this genus match those of the amended description of the genus as outlined by Saha et al. [32]. The genus belongs to the order *Xanthomonadales* and the family *Xanthomonadaceae*, based on the deep phylotaxonogenomic analysis in the present study.

### Emended description of the genus *Chiayiivirga* Hsu et al. 2013

Chi.a.yi.i.vir’ga. N.L. neut. n. *Chiayium*, Chiayi, a city in Taiwan, from where the type strain of the type species was isolated; L. fem. n. *virga*, stick; N.L. fem. n. *Chiayiivirga*, stick of Chiayi, a rod-shaped bacterium from Chiayi city. The type species of the genus is *Chiayiivirga flava* [31]. The genus *Chiayiivirga* belongs to order *Xanthomonadales* and family *Xanthomonadaceae*, not *Rhodanobacteraceae*, based on the deep phylotaxonogenomic analysis in the present study.

### Emended description of the genus *Rehaibacterium* Yu et al. 2013

Re.hai.bac.te’ri.um. N.L. n. *Rehaus*, Rehai referring to the isolation of the organism from Rehai National Park, Tengchong, Yunnan Province, south-west China; N.L. neut. n. *bacterium*, a small rod; N.L. neut. n. *Rehaibacterium*, a small rod from Rehai National Park. Type species of the genus is *Rehaibacterium terrae* [41]. The genus *Rehaibacterium* belongs to order *Xanthomonadales* and family *Xanthomonadaceae*, based on the deep phylotaxonogenomic analysis in the present study.

### Emended description of the genus *Pseudolysobacter* Wei et al. 2020

Pseu.do.ly.so.bac’ter. Gr. masc./fem. adj. *pseudês*, false; N.L. masc. n. *Lysobacter*, a bacterial genus; N.L. masc. n. *Pseudolysobacter*, a false *Lysobacter*. Type species of the genus *Pseudolysobacter* is *Pseudolysobacter antarcticus*. The genus *Pseudolysobacter* belongs to the family *Frateuriaceae* and the *order Xanthomonadales*, based on the deep phylotaxonogenomic analysis in the present study.

### Emended description of the genus Metallibacterium Ziegler et al. 2013

Me.tal.li.bac.te.ri.um. L. neut. n. metallum, mine; N.L. neut. n. bacterium, small rod; N.L. neut. n. Metallibacterium, a rod from a mine. Type species of the genus Metallibacterium is *Metallibacterium scheffleri* [36]. The genus *Metallibacterium* belongs to the family *Frateuriaceae* and the *order Xanthomonadales*, based on the deep phylotaxonogenomic analysis in the present study.

### Emended description of the genus *Mizugakiibacter* Kojima et al. 2014

Mi.zu.ga.ki.i.bac’ter. N.L. masc. n. bacter, a rod; N.L. masc. n. Mizugakiibacter, a rod isolated from Lake Mizugaki. Type species of the genus *Mizugakiibacter* is *Mizugakiibacter sediminis* [35]. The genus *Mizugakiibacter* belongs to the family *Frateuriaceae* and the *order Xanthomonadales*, based on the deep phylotaxonogenomic analysis in the present study.

### Emended description of the genus *Frateuria* Swing et al. 1980

Frat.eur’i.a. N.L. fem. n. Frateuria, named after Joseph Frateur (1903-1974), eminent Belgian microbiologist. Type species is *Frateuria aurantia* (ex Kondô and Ameyama 1958) Swings et al. 1980 [38]. *Frateuria* is a synonym of the previously described genera *Aerosticca, Fulvimonas, Rhodanobacter, Dyella, Luteibacter, Frateuria, Oleiagrimonas* and *Mizugakiibacter*. Description is as provided in Naushad et.al.2015 [1], and based on the deep phylotaxonogenomic analysis in the present study. The genus now includes genus *Aerosticca, Fulvimonas, Rhodanobacter, Dyella, Luteibacter, Frateuria, Oleiagrimonas* and *Mizugakiibacter*.

Emended descriptions of the member species for the genus *Frateuria* are provided in supplementary table 2.

### Emended description of the genus *Xanthomonas* Dowson 1939 (Approved Lists 1980)

Xan.tho.mo.nas. Gr. masc. adj. *xanthos*, yellow; L. fem. n. monas, unit, monad; N.L. fem. n. *Xanthomonas*, yellow monad.

The type genus is *Xanthomonas* (*Xanthomonas campestris* (Pammel 1895) Dowson 1939 (Approved Lists 1980))

Description is as provided in Naushad et.al., Saddler et.al. [1, 2], and based on the deep phylotaxonogenomic analysis in the present study. The genus now includes previously described genera i.e., *Xylella* [42], *Pseudoxanthomonas* [43], *Stenotrophomonas* [44], *Lysobacter* [45], *Vulcaniibacterium* [9], *Luteimonas* [43] and *Thermomonas* [46]. *Xanthomonas* is a synonym of the previously described genera *Xylella, Pseudoxanthomonas, Stenotrophomonas, Lysobacter, Vulcaniibacterium, Luteimonas*, and *Thermomonas*.

Emended description of the key member species for the genus *Xanthomonas* are provided below and remaining in the supplementary table 1.

**Emended description of *Xanthomonas maltophilia*** = *Stenotrophomonas maltophilia* ((Hugh 1981) Swings *et al*. 1983).

Description as provided in (Hugh 1981) Swings *et al*. 1983 and genomic analysis in the present study and a previous study by Bansal et.al. 2021 [24].

**Emended description of *Xanthomonas fastidiosa*** = *Xylella fastidiosa* Wells *et al*. 1987

Description as provided in Wells *et al*. 1987 and genomic analysis in the present study and a previous study by Bansal et.al., 2021 [24]

**Emended description of *Xanthomonas broegbernensis*** = *Pseudoxanthomonas broegbernensis* Finkmann *et al. 2000*

Description as provided in *Pseudoxanthomonas broegbernensis* Finkmann *et al*. 2000 and genomic analysis in the present study and a previous study by Bansal et.al., 2021 [24]

## Authors contribution

KB, SK and AS: Data analysis and drafting of manuscript, AC: Metadata preparation. PBP: Conceived and Participated in planning, design and drafting of the manuscript. All of the study. authors have read and approved the manuscript.

## Acknowledgement

This work was supported through NBRI-IMTECH-MLP48 and a council of scientific and industrial Research (CSIR) fellowship to AS.

## Transparency declaration

The authors declare that there are no conflicts of interest.

## Figure legends

**Supplementary Table 1**: List of member species of genus *Xanthomonas*

**Supplementary Table 2**: List of member species of genus *Frateuria*

**Supplementary Table 3:** Metadata showing total of 213 genomes belonging to type species and type strains of the order *Xanthomonadales*.

